# Disseminating cells in human oral tumours acquire an EMT cancer stem cell state that is predictive of metastasis

**DOI:** 10.1101/2020.04.07.029009

**Authors:** Gehad Youssef, Luke Gammon, Leah Ambler, Sophia Lunetto, Alice Scemama, Hannah Cottom, Kim Piper, Ian C. Mackenzie, Michael P. Philpott, Adrian Biddle

**Affiliations:** Blizard Institute, Barts and The London School of Medicine and Dentistry, Queen Mary University of London, UK; Department of Cellular Pathology, Barts Health NHS Trust, London, UK

## Abstract

Cancer stem cells (CSCs) undergo epithelial-mesenchymal transition (EMT) to drive metastatic dissemination in experimental cancer models. However, tumour cells undergoing EMT have not been observed disseminating into the tissue surrounding human tumour specimens, leaving the relevance to human cancer uncertain. We have previously identified both EpCAM and CD24 as markers of EMT CSCs with enhanced plasticity. This afforded the opportunity to investigate whether retention of EpCAM and CD24 alongside upregulation of the EMT marker Vimentin can identify disseminating EMT CSCs in human tumours. Examining disseminating tumour cells in over 12,000 imaging fields from 84 human oral cancer specimens, we see a significant enrichment of single EpCAM, CD24 and Vimentin co-stained cells disseminating beyond the tumour body in metastatic specimens. Through training an artificial neural network, these predict metastasis with high accuracy (cross-validated accuracy of 87-89%). In this study, we have observed single disseminating EMT CSCs in human oral cancer specimens, and these are highly predictive of metastatic disease.

## Introduction

In multiple types of carcinoma, cancer stem cells (CSCs) undergo epithelial-mesenchymal transition (EMT) to enable metastatic dissemination from the primary tumour (Biddle et al., 2011; Lawson et al., 2015; Liu et al., 2014; Ruscetti et al., 2016). This model of metastatic dissemination has been built from studies using murine models and human cancer cell line models. However, this process has not been observed in human tumours in the *in vivo* setting, leading to uncertainty over the relevance of these findings to human tumour metastasis (Bill and Christofori, 2015; Williams et al., 2019). A key complication with efforts to study metastatic processes in human tumours is the inability to trace cell lineage. As cancer cells exiting the tumour downregulate epithelial markers whilst undergoing EMT, they become indistinguishable from the mesenchymal non-tumour cells surrounding the tumour (Li and Kang, 2016). Therefore, once these cells detach from the tumour body and move away they are lost to analysis. Attempts have been made to use the retention of epithelial markers alongside acquisition of mesenchymal markers to identify cells undergoing EMT in human tumours (Bronsert et al., 2014; Jensen et al., 2015; Puram et al., 2017). However, these studies were limited to characterising cells undergoing the earliest stages of EMT whilst still attached to the cohesive body of the primary tumour.

EMT must be followed by the reverse process of mesenchymal-to-epithelial transition (MET) to enable new tumour growth at secondary sites, and therefore retained plasticity manifested as ability to revert to an epithelial phenotype is an important feature of metastatic CSCs (Ocana et al., 2012; Tsai et al., 2012). We have previously demonstrated that a CD44^high^EpCAM^low/-^ EMT population can be separated from the main CD44^low^EpCAM^high^ epithelial population in flow cytometric analysis of oral squamous cell carcinoma (OSCC) cell lines and fresh tumour specimens (Biddle et al., 2016; Biddle et al., 2011). We identified retained cell surface expression of EpCAM (Biddle et al., 2011) and CD24 (Biddle et al., 2016) in a minority of cells that have undergone a full morphological EMT. Both EpCAM and CD24 were individually associated with enhanced ability to undergo MET, and thus are markers of EMT CSCs exhibiting retained plasticity. We therefore reasoned that retention of one or both of these markers may identify an important population of tumour cells that have undergone EMT and disseminated from the primary tumour in human tumour specimens, and are responsible for subsequent metastatic seeding. Here, we characterise the combined role of EpCAM and CD24 in marking a population of disseminating tumour cells in human OSCC specimens. Staining for EpCAM and CD24 alongside the mesenchymal marker Vimentin in over 12,000 imaging fields from 84 human tumour specimens, stratified on metastatic status, identifies cells that have undergone EMT and disseminated into the stromal region surrounding metastatic primary tumours. Using a machine learning approach, we show that the presence of these EMT CSCs in the tumour stroma is predictive of metastasis.

## Results

### Identification of human tumour cells that have undergone an EMT and disseminated into the surrounding stromal region

The retention of EpCAM expression in a sub-population of tumour cells that have undergone EMT raised the prospect that we may be able to identify these cells outside of the tumour body in human tumour specimens, as EpCAM is a specific epithelial marker that would not normally be found in the surrounding stromal region. In combination with EpCAM, we stained tumour specimens for CD24 as a second marker of plastic EMT CSCs, and Vimentin as a mesenchymal marker to identify cells that have undergone EMT. Notably, CD44 cannot be used as an EMT marker in the context of intact tissue as it requires trypsin degradation in order to yield differential expression in EMT and epithelial populations (Biddle et al., 2013; Mack and Gires, 2008). Vimentin, on the other hand, accurately distinguishes EMT from epithelial tumour cells in immunofluorescent staining protocols (Biddle et al., 2016). By combining EpCAM as a tumour lineage and EMT CSC marker, Vimentin as a mesenchymal marker, and CD24 as a plastic EMT CSC marker, we aimed to identify tumour cells that have undergone EMT and disseminated into the surrounding stromal region. For this, we developed a protocol for automated 4-colour (3 markers + nuclear stain) immunofluorescent imaging and analysis of entire histopathological slide specimens, to test for co-localisation of the 3 markers in each individual cell across each specimen.

To determine whether this marker combination identifies EMT CSCs, we initially tested the protocol on the CA1 OSCC cell line and an EMT CSC sub-line that is a derivative of this cell line (EMT-stem sub-line) (Biddle et al., 2016). EpCAM^+^Vim^+^CD24^+^ cells were greatly enriched in the EMT-stem sub-line, comprising 41% of the population, compared to 2.1% in the CA1 line (Figure 1A, B, E). Cells with this staining profile were absent from normal keratinocyte culture and cancer associated fibroblast culture (Supplementary Figure S1). To test the specific role of EpCAM retention, we replaced EpCAM with a pan-keratin antibody against epithelial keratins. There was very little Pan-keratin^+^Vim^+^CD24^+^ staining, and no enrichment for Pan-keratin^+^Vim^+^CD24^+^ cells in the EMT-stem sub-line (Figure 1C, D, E). Therefore, whilst epithelial keratins are lost, EpCAM is retained in cells undergoing EMT and an EpCAM^+^Vim^+^CD24^+^ staining profile can be used as a marker for EMT CSCs in immunofluorescent staining protocols.

**Figure 1.**
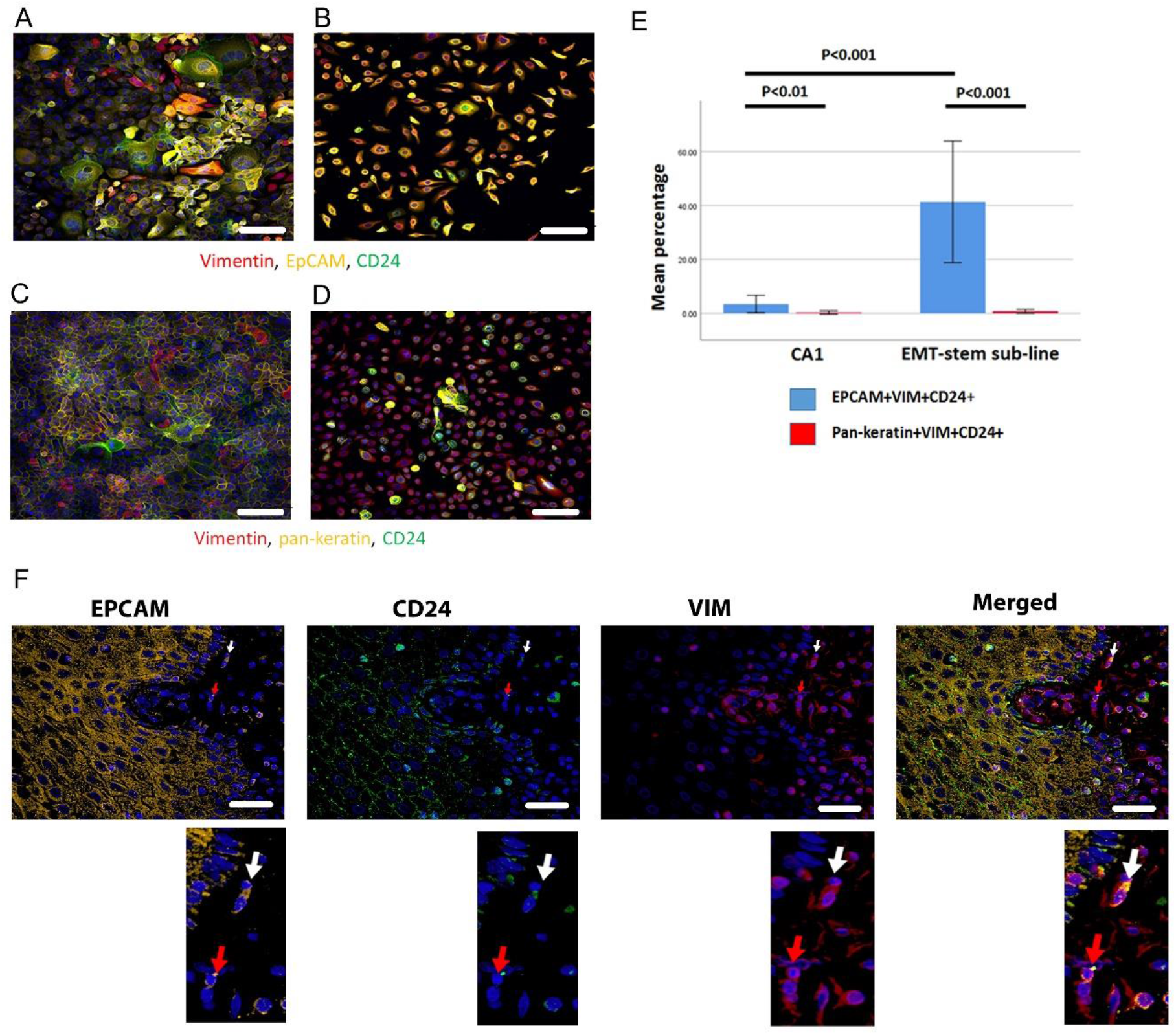
Immunofluorescent co-staining for EpCAM, Vimentin and CD24 identifies the EMT stem cell state. **A-D**, Immunofluorescent staining for EpCAM, Vimentin and CD24 (A, B) and pan-keratin, Vimentin and CD24 (C, D) in the CA1 cell line (A, C) and the EMT-stem CA1 sub-line (B, D). **E**, Quantification of the percentage of EpCAM^+^Vim^+^CD24^+^ and pan-keratin^+^Vim^+^CD24^+^ cells in the CA1 cell line and EMT-stem sub-line. Significance is obtained from a two-tailed student t-test. The graph shows mean +/-95% confidence interval. **F**, Detection of EpCAM^+^Vim^+^CD24^+^ cells in the stroma surrounding an oral cancer tumour specimen. The white arrow highlights an EpCAM^+^Vim^+^CD24^+^ cell in the stroma. The red arrow highlights an EpCAM^+^Vim^+^CD24^-^ cell in the stroma. DAPI nuclear stain is blue. Below inset; enlargement of the highlighted cells for each marker. Scale bars = 100µm.

Imaging the tumour body and adjacent stroma in sections of human OSCC specimens, we detected single cells co-expressing EpCAM, Vimentin and CD24 in the stromal region surrounding the tumour (Figure 1F), confirming that these cells can be detected in human tumour specimens. We next stratified 24 human primary OSCC specimens into 12 tumours that had evidence of lymph node metastasis or perineural spread, and 12 that remained metastasis free (Supplementary Figure S2), and stained them for EpCAM, Vimentin and CD24. Single cells co-expressing EpCAM, Vimentin and CD24 were abundant in the stroma surrounding metastatic tumours. This was not the case in non-metastatic tumours or normal epithelial regions (Figure 2, A-C). In contrast to EpCAM, pan-keratin staining did not identify cells in the stroma surrounding metastatic tumours (Figure 2D).

**Figure 2.**
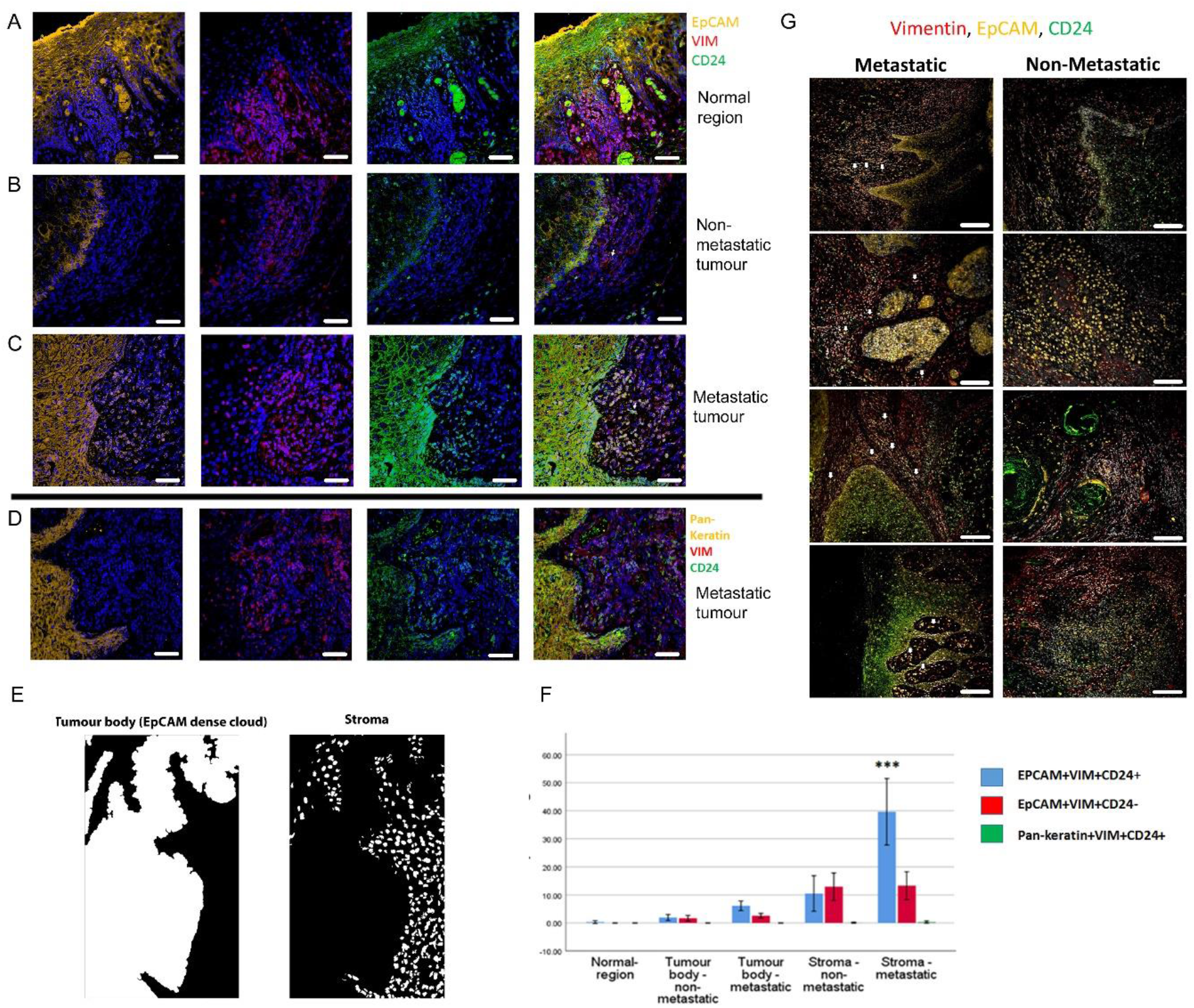
Enrichment of EpCAM^+^Vim^+^CD24^+^ cells in the stroma surrounding metastatic tumours. **A-C**, Immunofluorescent four-colour staining of oral tumour specimens for EpCAM (yellow), Vimentin (red) and CD24 (green) with DAPI nuclear stain (blue). Representative imaging fields from a normal epithelial region (A), a non-metastatic tumour (B) and a metastatic tumour (C). **D**, Staining of a metastatic tumour for pan-keratin, Vimentin and CD24. **E**, Image segmentation was performed, with generation of an ‘EpCAM dense cloud’ to distinguish the tumour body from the stroma. Grey level intensities for EpCAM, Vimentin and CD24 were obtained for every nucleated cell in each imaging field. **F**, Quantification of the percentage of EpCAM^+^Vim^+^CD24^+^, EpCAM^+^Vim^+^CD24^-^ and pan-keratin^+^Vim^+^CD24^+^ cells in normal region (epithelium distant from the tumour), tumour body, and stromal region from metastatic and non-metastatic tumours in the first batch of specimens. A student t-test was performed comparing the mean percentage of EpCAM^+^Vim^+^CD24^+^ co-expressing cells in the metastatic stroma compared to the other fractions. *** signifies p < 0.001. The graph shows mean +/-95% confidence interval. **G**, Immunofluorescent four-colour staining of oral tumours from the second batch of specimens, showing tumours with a range of invasive front presentations. White arrows highlight single EpCAM^+^Vim^+^CD24^+^ cells in the stroma. Scale bars = 100 µm.

**Figure 3.**
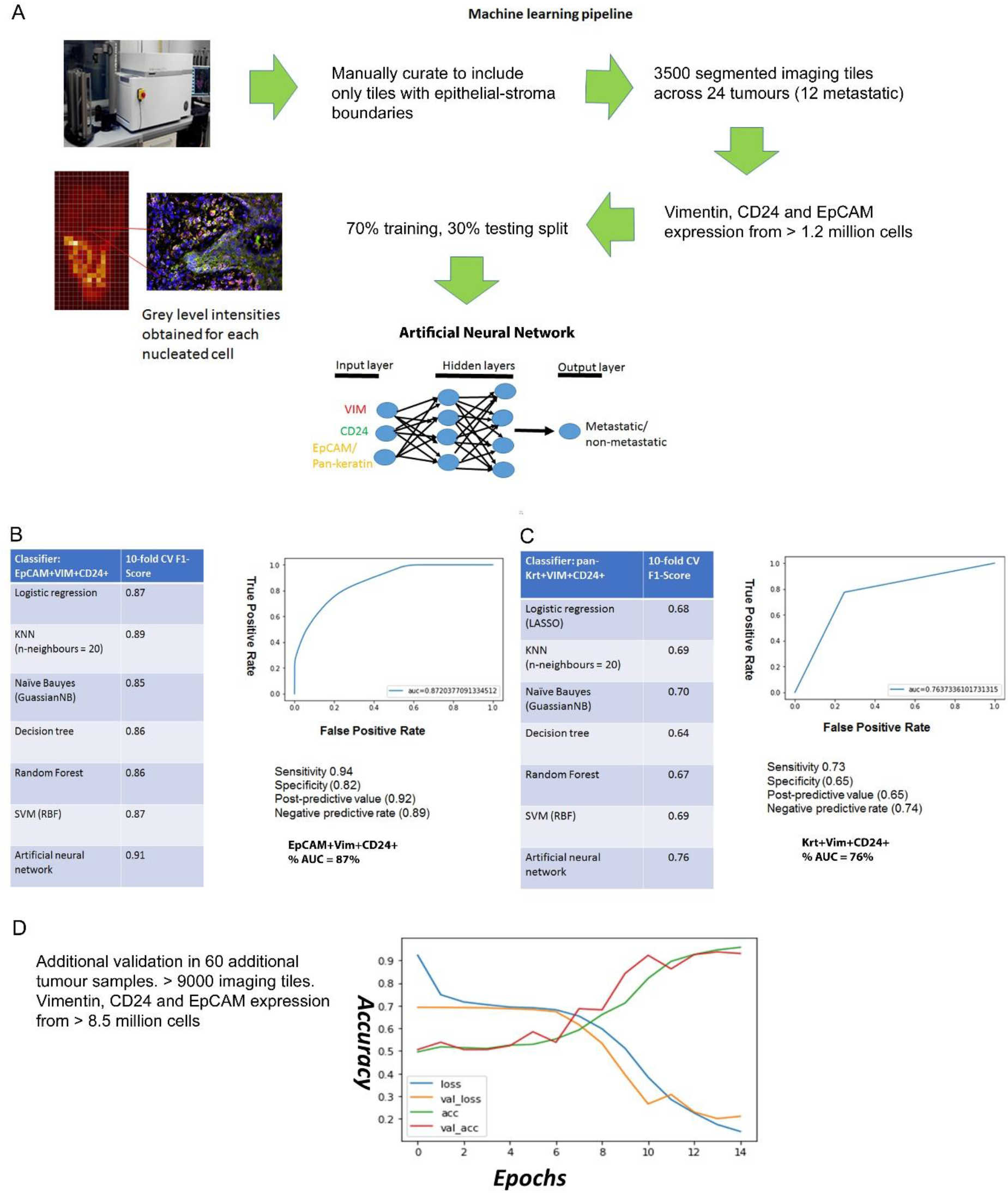
Predicting metastasis using EpCAM, Vimentin and CD24 immunofluorescent staining and a supervised machine learning approach. **A**, Pipeline for machine learning based on grey level intensities for the three markers in tumour batch 1. The training tiles were classified as coming from a metastatic or non-metastatic tumour. **B, C**, Performance of EpCAM, Vimentin and CD24 (B) and pan-keratin, Vimentin and CD24 (C) in the supervised learning task on tumour batch 1. The tables show the 10-fold cross-validation F1 scores of different machine learning classification algorithms. To the right of each table is a receiver-of-operator curve (ROC) showing the area under the curve (AUC) of the artificial neural network (ANN) classifier. **D**, Performance of EpCAM, Vimentin and CD24 in the supervised learning task on tumour batch 2. An ANN classifier was trained and tested on batch 2, independently of tumour batch 1. Accuracy and loss scores are displayed for the training set (green and blue lines) and the validation set (red and yellow lines) drawn from within this batch, for 14 training epochs on the ANN classifier.

We developed an image segmentation protocol that separated the tumour body from the adjacent stroma, thus enabling each nucleated cell to be assigned to either the tumour or stromal region in automated image analysis (Figure 2E). Expression of EpCAM, Vimentin and CD24 was then analysed for every nucleated cell in every imaging field that included both tumour and stroma (3500 manually curated imaging fields across the 24 tumours). This enabled the proportion of each cell type in each region to be quantified (Figure 2F). EpCAM^+^Vim^+^CD24^+^ cells were enriched in the stroma compared to the tumour body, and there was a much greater accumulation of EpCAM^+^Vim^+^CD24^+^ cells in the stroma of metastatic tumours compared to non-metastatic tumours. Interestingly, this was not the case for EpCAM^+^Vim^+^CD24^-^ cells, which were also enriched in the stroma but showed no difference between metastatic and non-metastatic tumours. Pan-keratin^+^Vim^+^CD24^+^ cells were not detected.

To extend this analysis, we stained and imaged a further 60 tumours, evenly stratified on the same criteria. These displayed the same evidence of individual disseminating cells co-expressing EpCAM, Vimentin and CD24 in metastatic tumours only (Figure 2G and Supplementary Figure S3F, G). For these tumours, using a variation on the previous image segmentation protocol (Supplementary Figure S3, A-D), the proportion of EpCAM^+^Vim^+^CD24^+^ and EpCAM^+^Vim^+^CD24^-^ cells was quantified for each cell in over 9000 imaging fields at the tumour-stroma boundary (Supplementary Figure S3E). Consistent with the previous set of tumours, only EpCAM^+^Vim^+^CD24^+^ cells were specifically enriched in the stroma of metastatic tumours.

To explore whether these EpCAM^+^Vim^+^CD24^+^ cells in the stroma may in fact be non-tumour cell types, we analysed a published scRNAseq dataset for human head and neck cancer (Puram et al., 2017). In this dataset, tumour and non-tumour cells were separated using bioinformatic techniques (principally inferred CNV and a ‘tumour-epithelial’ expression signature). Analysing this dataset for EpCAM, Vimentin and CD24 co-expression, we found that 12% of tumour cells (267/2215) were EpCAM^+^Vim^+^CD24^+^. In the non-tumour cells, only 0.8% (29/3687) were EpCAM^+^Vim^+^CD24^+^ (Supplementary Figure S4). Therefore, the observed EpCAM^+^Vim^+^CD24^+^ cells in our tumour specimens are highly likely to be a tumour cell population. Indeed, use of EpCAM as a tumour lineage marker is specifically intended to exclude staining for stromal constituents. EpCAM is a specific epithelial marker, that is not expressed in stromal or immune cells – it is expressed exclusively in epithelia and epithelial-derived tumours (Keller et al., 2019).

These findings demonstrate that an EpCAM^+^Vim^+^CD24^+^ staining profile marks tumour cells disseminating into the surrounding stroma, and that these cells are enriched specifically in metastatic tumours. The presence of disseminating tumour cells that express EpCAM but not CD24 did not correlate with metastasis. This highlights a requirement for the plasticity marker CD24, when identifying disseminating metastatic CSCs.

### Identification of EpCAM^+^CD24^+^Vim^+^ CSCs enables clinical prediction using a machine learning approach

OSCC are an important health burden and one of the top ten cancers worldwide, with over 300,000 cases annually and a 50% 5-year survival rate. There is frequent metastatic spread to the lymph nodes of the neck; this is the single most important predictor of outcome and an important factor in treatment decisions (Sano and Myers, 2007). If spread to the lymph nodes is suspected, OSCC resection is accompanied by neck dissection to remove the draining lymph nodes, a procedure with significant morbidity. At presentation it is currently very difficult to determine which tumours are metastatic and this results in sub-optimal tailoring of treatment decisions. Accurate prediction of metastasis would therefore have great potential to improve clinical management of the disease to reduce both mortality and treatment-related morbidity. We sought to determine whether the EpCAM^+^CD24^+^Vim^+^ staining pattern could be predictive of metastasis.

Starting with the EpCAM, Vimentin and CD24 immunofluorescence grey levels for each nucleated cell, we used a supervised machine learning approach to predict whether an imaging field comes from a metastatic or non-metastatic tumour (Figure 5A). As a benchmark we used the pan-keratin, Vimentin and CD24 immunofluorescence grey levels, as we hypothesised that pan-keratin would provide an inferior predictive value than EpCAM given that there was no dissemination of pan-keratin expressing cells in the stroma. 3500 imaging fields containing 2,640,000 total nucleated cells from 24 tumour specimens were used in the machine learning task. We compared the performance accuracy (10-fold cross-validated F-score) of different machine learning classification algorithms. The best performing classifiers for EpCAM, Vimentin and CD24 were the artificial neural network (ANN) and support vector machine (SVM), with F1 accuracy scores of 91% and 87% respectfully (Figure 5B). For the ANN, the area under the curve (AUC) accuracy score was 87%, with 94% sensitivity and 82% specificity. Training with Pan-keratin, Vimentin and CD24 gave much worse prediction across all classifiers (Figure 5C). These findings demonstrate that, utilising a machine learning algorithm, staining for EpCAM, Vimentin and CD24 can predict metastatic status with high accuracy and may therefore have clinical utility.

To extend this analysis of utility for metastasis prediction, we stained and imaged a further 60 tumours, evenly stratified on the same criteria, for EpCAM, Vimentin and CD24. Over 9000 imaging fields at the tumour-stroma boundary from 60 evenly stratified tumour specimens, containing over 8.5 million nucleated cells, were fed into an artificial neural network machine learning task. For this task, we recorded the predictive accuracy from the training and validation sets after each training epoch, which showed good alignment and an 89% accuracy score after 12 training epochs (Figure 5D).

To our knowledge, this is the first time immunofluorescent staining of human tumour tissue specimens has been used in a machine learning pipeline for clinical prediction. Previous studies using cytokeratin immunohistochemistry, clinicopathological data and serum biomarkers for clinical prediction via machine learning have achieved AUCs of 75% in breast cancer (Tseng et al., 2019), 80% in OSCC (Bur et al., 2019), and 82% in colorectal cancer (Takamatsu et al., 2019).

## Discussion

The role of EMT in tumour dissemination has long been debated but, lacking evidence of cells undergoing EMT whilst disseminating from human tumours *in vivo*, this role has had to be inferred from mouse models and human cell line models. Here, through applying our understanding of EMT cancer cell heterogeneity and markers for plastic EMT CSCs, we have identified EMT CSCs disseminating from the primary tumour in human pathological specimens. Importantly, the presence of these disseminating stem cells is strongly correlated with tumour metastasis. Using a machine learning approach, we have demonstrated the ability to predict metastasis with high accuracy through staining for these EMT CSCs.

A partial EMT state has previously been identified in an OSCC scRNAseq dataset; this state retained epithelial gene expression alongside expression of mesenchymal genes, and was correlated with nodal metastasis and adverse pathological features (Puram et al., 2017). Here, using immunofluorescent staining for EMT CSCs that retain the epithelial marker EpCAM alongside the mesenchymal marker Vimentin and the CSC plasticity marker CD24, we have identified single EMT CSCs disseminating into the stroma surrounding oral tumours. However, epithelial keratins are not retained. We have also shown that retention of EpCAM is not on its own sufficient alongside Vimentin to mark disseminating EMT CSCs that correlate with metastasis. There is a requirement for CD24, which we have previously shown to be a plasticity marker within the EMT population even when driven into full morphological EMT under TGFβ treatment (Biddle et al., 2016). This suggests that the EMT CSC state may be more complex than a simple coalescence of epithelial and mesenchymal characteristics.

We have identified an EMT CSC state that disseminates as single cells from human tumours and is correlated with metastasis. Immunofluorescent antibody co-staining for EpCAM, CD24 and Vimentin identifies these EMT CSCs in human tumour specimens and is predictive of metastasis. The ability of this co-staining to separate disseminating tumour cells from the stromal content of human tumours, which has confounded previous attempts to develop a predictive EMT signature (Tan et al., 2014), is one important factor in this success. However, we also show that EpCAM^+^CD24^-^Vim^+^ tumour cells in the stroma do not correlate with metastasis, and therefore the clinically predictive utility of tumour cell staining in the stroma can be isolated specifically to the EpCAM^+^CD24^+^Vim^+^ EMT CSCs. This highlights the value of using techniques that give single cell resolution, enabling isolation of the signal to the specific cell type of interest within a highly heterogeneous cellular environment. An important strength of our study has been the ability to look at the single cell level in an automated fashion across thousands of fields of view from human tumours, enabling us to observe and quantify human tumour cells disseminating into the surrounding tissue. In doing so, we have identified single disseminating EMT CSCs that are predictive of metastasis.

## Supporting information

Supplementary information for Youssef et al

## Conflict of interest

The authors declare no conflicts of interest.

## Acknowledgements

We thank Ryan O’Shaughnessy, Sarah Marzi, and Jan Soetaert for technical assistance and discussion. Gehad Youssef and Adrian Biddle were supported by Animal Free Research UK, as part of the Animal Replacement Centre of Excellence at Queen Mary University of London. Leah Ambler was supported by Oracle Cancer Trust.

## Methods

### Cell culture

The CA1 OSCC cell line and oral cancer associated fibroblasts were both previously derived in our laboratory, from separate biopsies of OSCC of the floor of the mouth. The EMT-stem sub-line was derived as a single cell clone from the CA1 cell line (Biddle et al., 2016). Normal keratinocytes were the N/TERT hTERT-immortalised epidermal keratinocyte cell line (Smits et al., 2017). Cell culture was performed as previously described (Biddle et al., 2011). Cell removal from adherent culture was performed using 1x Trypsin-EDTA (Sigma, T3924) at 37^º^C.

### Immunofluorescent staining of cell lines and tumour tissue sections

Tumour specimens were obtained from the pathology department at Barts Health NHS Trust, with full local ethical approval and patients’ informed consent. Sections of formalin fixed paraffin embedded (FFPE) archival specimens were dewaxed by clearing twice in xylene for 5 minutes then gradually hydrating the specimens in an alcohol gradient (100%, 90%, 70%) for 3 minutes each. The sections were then washed under running tap water before immersing the slides in Tris-EDTA pH9 for antigen retrieval using a standard microwave at high power for 2 minutes and then 8 minutes at low power.

Four-colour immunofluorescent staining was performed by firstly staining the membranous proteins prior to the permeabilisation and blocking steps. The sections were incubated with an IgG2a mouse monoclonal CD24 antibody (clone ML5, BD Bioscience) and IgG rabbit recombinant monoclonal EpCAM antibody (EPR20532-225, Abcam) in PBS overnight at 4°C (1/100 dilution). The sections were then washed three times in PBS and incubated for 1 hour at room temperature with anti-mouse IgG2 Alexa Fluor 488 and anti-rabbit IgG Alexa Fluor 555 secondary antibodies (1/500 dilution). The sections were then washed in PBS and permeabilised with 0.5% triton-X in PBS for 10 minutes followed by blocking for 1 hour with blocking buffer (3% goat serum, 2% bovine serum albumin in PBS). The sections were then incubated with an IgG1 mouse monoclonal Vimentin antibody (clone V9, Dako) and (optionally, in place of EpCAM) IgG rabbit polyclonal wide spectrum cytokeratin antibody (ab9377, Abcam) overnight at 4°C in blocking buffer (1/100 dilution). After washing with PBS, the sections were incubated with anti-mouse IgG1 Alexa Fluor 647 antibody and (optionally) anti-rabbit IgG Alexa Fluor 555 for 1hr at 4°C (1/500 dilution). After washing three times with PBS, cell nuclei were stained with DAPI (1/1000 dilution in PBS) for 10 minutes.

For cell line staining, cells were fixed in 4% PFA for 10 minutes then washed with PBS. Staining was performed in the same manner as described above, however permeabilisation was performed with 0.25% Triton-X for 10 minutes and DAPI incubation was reduced to 1 minute.

### Quantifying the abundance of stained sub-populations in cell lines and tumour tissue sections

Imaging of the stained slides was performed using the In Cell Analyzer 2200 (GE), a high content automated fluorescence microscope with four-colour imaging capability. The slides were imaged at x20 and x40 magnification. An image segmentation protocol was developed to extract grey level intensities corresponding to EpCAM, Vimentin and CD24 expression for every DAPI stained nucleated cell in the tumour body and the adjacent stroma separately. Segmentation was performed using the Developer Toolbook software (GE). As shown in figure 2E and Supplementary Figure S3, an ‘EpCAM dense cloud’ or ‘Vimentin dense cloud’ was generated to isolate individual nucleated cells in the tumour body from the adjacent stroma and analyse them separately.

Grey level intensities obtained from the imaging analysis were processed in the following way. Firstly, the median number of nucleated cells was calculated and imaging fields with fewer than 20% of the median nucleated cells were excluded from the analysis pipeline. The folded edges of a specimen were also excluded. The median grey level intensity of the FITC, CY3 and CY5 fluorescence channels corresponding to CD24, EpCAM and Vimentin expression were computed for the negative control stained slides. A nucleated cell was deemed to have positive CD24, EpCAM or Vimentin expression if its grey level intensity exceeded the background threshold value (1.5 x median grey level intensity of negative control slide) for the FITC, CY3 and CY5 channels respectively. If a nucleated cell surpassed the background threshold for all three fluorescence channels it was termed a triple positive cell (CD24^+^EpCAM^+^Vim^+^) and denoted with 1 and if this criteria was not met the nucleated cell was denoted with a 0. For EpCAM^+^Vim^+^CD24^-^ cells (termed double positive), the nucleated cell must exceed the background threshold for the CY3 and CY5 channels but not the FITC.

The scRNAseq dataset (Puram et al., 2017) was analysed using a threshold (median or quartile) using the normalised count expression for EpCAM, CD24 and Vimentin for each cell.

### Machine learning for prognostic prediction using immunofluorescent staining data

A dataset was created of a pool of 2,640,000 nucleated cells across 3500 imaging fields from 24 tumour specimens (12 with lymph node metastasis or perineural spread, and 12 without) (batch 1) or 8,563,000 nucleated cells across 9,200 imaging fields from 60 tumour specimens (30 with lymph node metastasis or perineural spread, and 30 without) (batch 2). The background threshold for the FITC, CY3 and CY5 channels was subtracted from the grey level intensities for each nucleated cell. The supervised machine learning task was to classify each imaging field into whether it belonged to a metastatic or non-metastatic tumour.

The dataset was stratified into a training and validation cohort in a 70%:30% ratio using a random seed split. Supervised machine learning approaches were implemented using the skikit-learn Python 3.6 libraries (Pedregosa et al., 2011) and Tensorflow/Keras framework (https://www.tensorflow.org/api_docs/python/tf/keras/models). Hyper-parameter optimisation was performed by an exhaustive grid search and computed on Apocrita, a high performance cluster (HPC) facility at Queen Mary University of London (http://doi.org/10.5281/zenodo.438045). To further minimise overfitting, 10-fold cross-validation was performed and the mean accuracy metric, F1 score, was obtained for each learning iteration. Receiver-of-operator (ROC) curves and the area-under the-curve (AUC) were computed for the optimum supervised learning algorithm. Supervised approaches used were logistic regression, support vector machines (Smola and Scholkopf, 2004), Naïve Bayes (Zhang, 2005), K-Nearest Neighbours (Bentley, 1975), decision trees (Dumont et al., 2009), and artificial neural networks (Rumelhart et al., 1986).

## Notes

### Competing Interest Statement

The authors have declared no competing interest.

### Summary of Updates

In response to the attached referee comments, we have substantially revised this manuscript. We have also streamlined it to focus on the central finding; that we can detect individual disseminating cancer stem cells in human tissue and these are predictive of metastatic progression.

