## Supplementary information for Youssef et al for "Disseminating cells in human oral tumours acquire an EMT cancer stem cell state that is predictive of metastasis"

**Supplementary Figure S1**

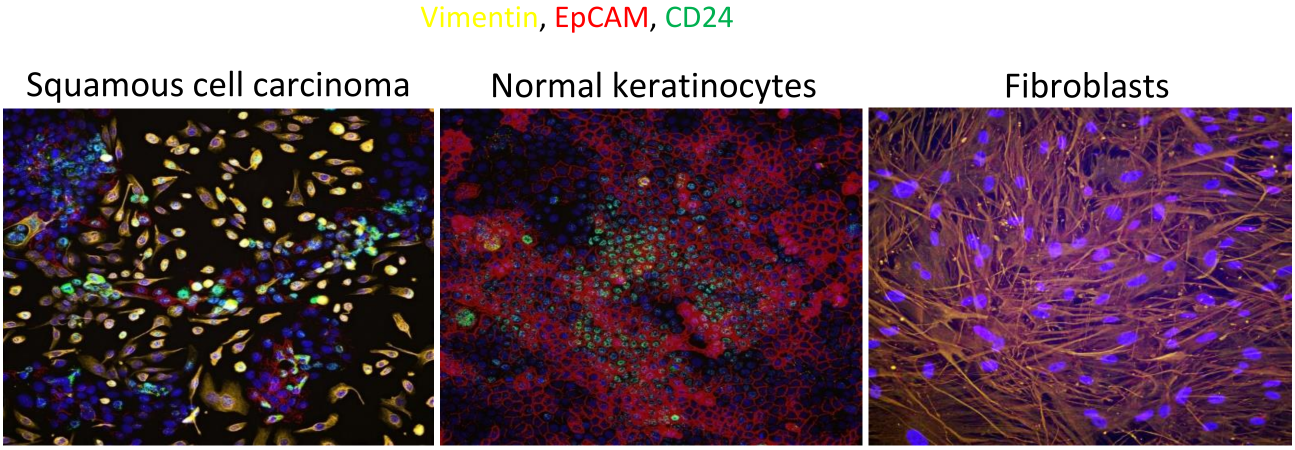


Figure S1 – EpCAM, Vimentin and CD24 immunofluorescent staining in the CA1 OSCC cell line (left), normal keratinocytes (centre) and oral cancer associate fibroblasts (right). Yellow = Vimentin, Red = EpCAM, Green = CD24.

**Supplementary Figure S2**

| Tumour | Nodal metastatic | Tumour site | perineural spread | Depth of invasion | Tumour stage [TNM 7th Ed] | Tumour Differentiation | Pattern of invasion |
|---|---|---|---|---|---|---|---|
| 1 | yes | Left lateral tongue | yes | 11mm | pT3 pN2b | Moderate to poorly differentiated | dis-cohesive |
| 2 | yes | Right tongue / floor of mouth | No | 10mm | pT3 pN2b | Moderately differentiated | dis-cohesive |
| 3 | yes | Maxillary sulcus | No | 8.5mm | pT2 pNx | Moderately differentiated | dis-cohesive |
| 4 | yes | Right floor of mouth | No | 3mm | pT1 pNx | Moderate to poorly differentiated | dis-cohesive |
| 5 | yes | Left posterior ventro-lateral tongue and left tonsillar fossa | Yes | 7.2mm | pT4a pN2C | Moderate to poorly differentiated | dis-cohesive |
| 6 | yes | Left lateral border of tongue | Yes | 21mm | pT2 pN2b | Moderate to poorly differentiated | dis-cohesive |
| 7 | yes | Left tongue | Yes | 12mm | pT2 pN2b | Moderate to poorly differentiated | dis-cohesive |
| 8 | yes | Right buccal mucosa | Yes | 10mm | pT2 pN2b | Moderate to poorly differentiated | dis-cohesive |
| 9 | yes | Left tongue | Yes | 13.5mm | pT2 pN2b | Poorly differentiated | dis-cohesive |
| 10 | yes | Left buccal mucosa | yes | 11.5mm | pT2 pN2b | Moderate and focally poorly differentiated | cohesive |
| 11 | yes | Left tongue | yes | 11.5mm | pT2 pN2b | Moderate to poorly differentiated | dis-cohesive |
| 12 | No | Left tongue | yes | 7.9mm | pT2 pN0 | Poorly differentiated | dis-cohesive |
| 13 | No | Left buccal mucosa | No | 10mm | pT1 pNx, | Well to moderately differentiated | cohesive |
| 14 | No | Right tongue | No | 7.5mm | pT1 pN0 | Moderately differentiated | dis-cohesive |
| 15 | No | Left tongue | No | 5mm | pT1 pN0 | Moderately differentiated | dis-cohesive |
| 16 | No | Left tongue | No | 2.5mm | pT1 | Moderately differentiated | cohesive |
| 17 | No | Right soft palate | No | 6.5mm | pT1 | Moderate to poorly differentiated | dis-cohesive |
| 18 | No | Midline anterior mandibular gingiva | No | 4mm | pT4a pN0 | Well to moderately differentiated | dis-cohesive |
| 19 | No | Left mandibular alveolus | No | 13.5mm | pT4a pN0 | Moderately differentiated | dis-cohesive |
| 20 | No | Left dorsum of tongue | No | 1.5mm | pT1 | Well differentiated | cohesive |
| 21 | No | Right anterior tongue | No | 5mm | pT2 pNx | Moderate to poorly differentiated | cohesive |
| 22 | No | Left floor of mouth / ventral tongue | No | 3mm | pT1 | Well to moderately differentiated | cohesive |
| 23 | No | Right buccal mucosa | No | 3.5mm | pT1 | Moderately differentiated | cohesive |
| 24 | No | Lower right labial mucosa / vermilion | No | 3.5mm | pT1 | Moderately differentiated | dis-cohesive |

Figure S2 – Tumour characteristics for the 12 metastatic tumours (tumours 1 – 12) and 12 non-metastatic tumours (tumours 13 – 24) that were sectioned and stained for EpCAM, CD24 and Vimentin in the first batch.

**Supplementary Figure S3**

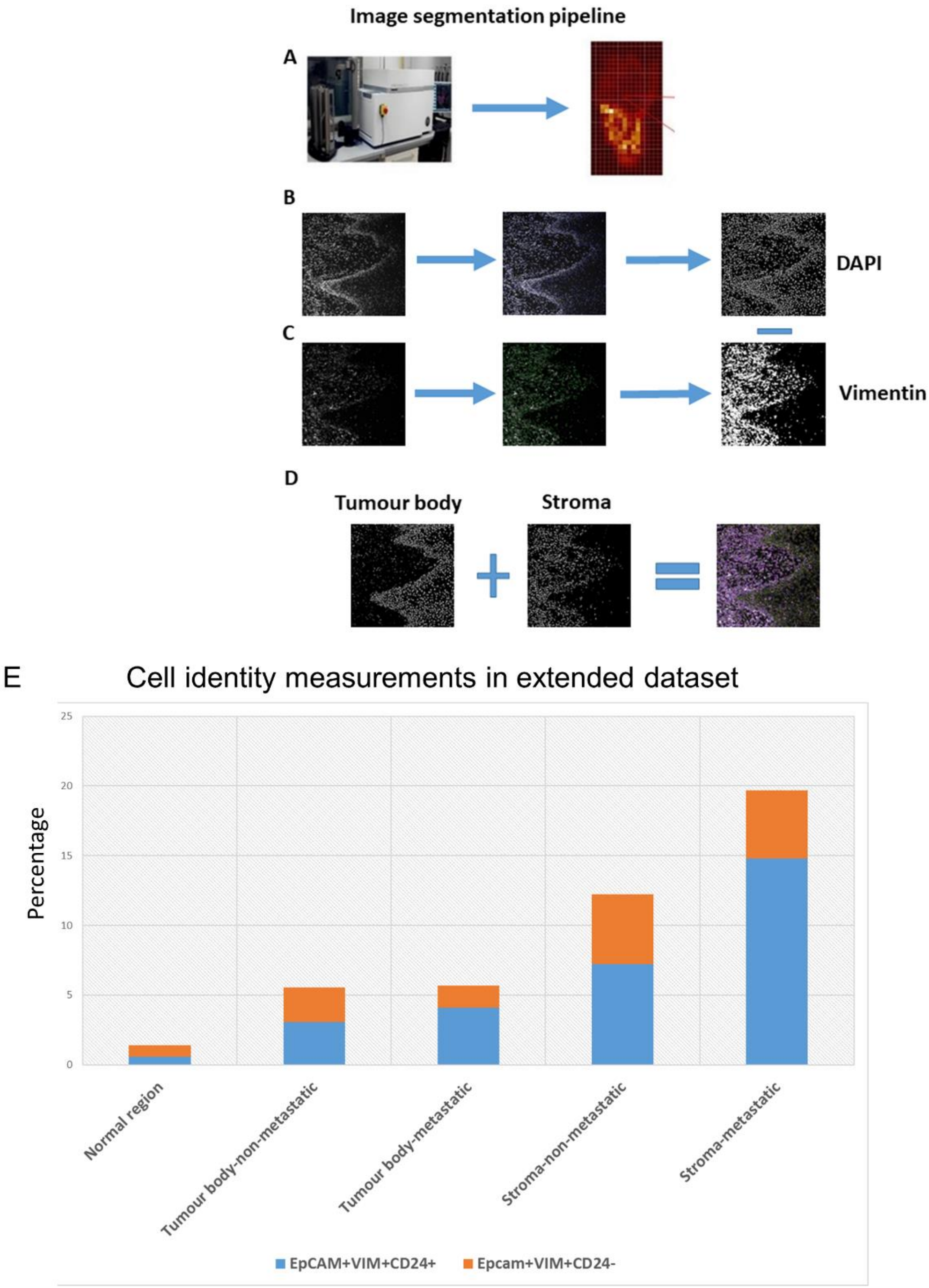

F Metastatic tumours Vim, EpCAM, CD24

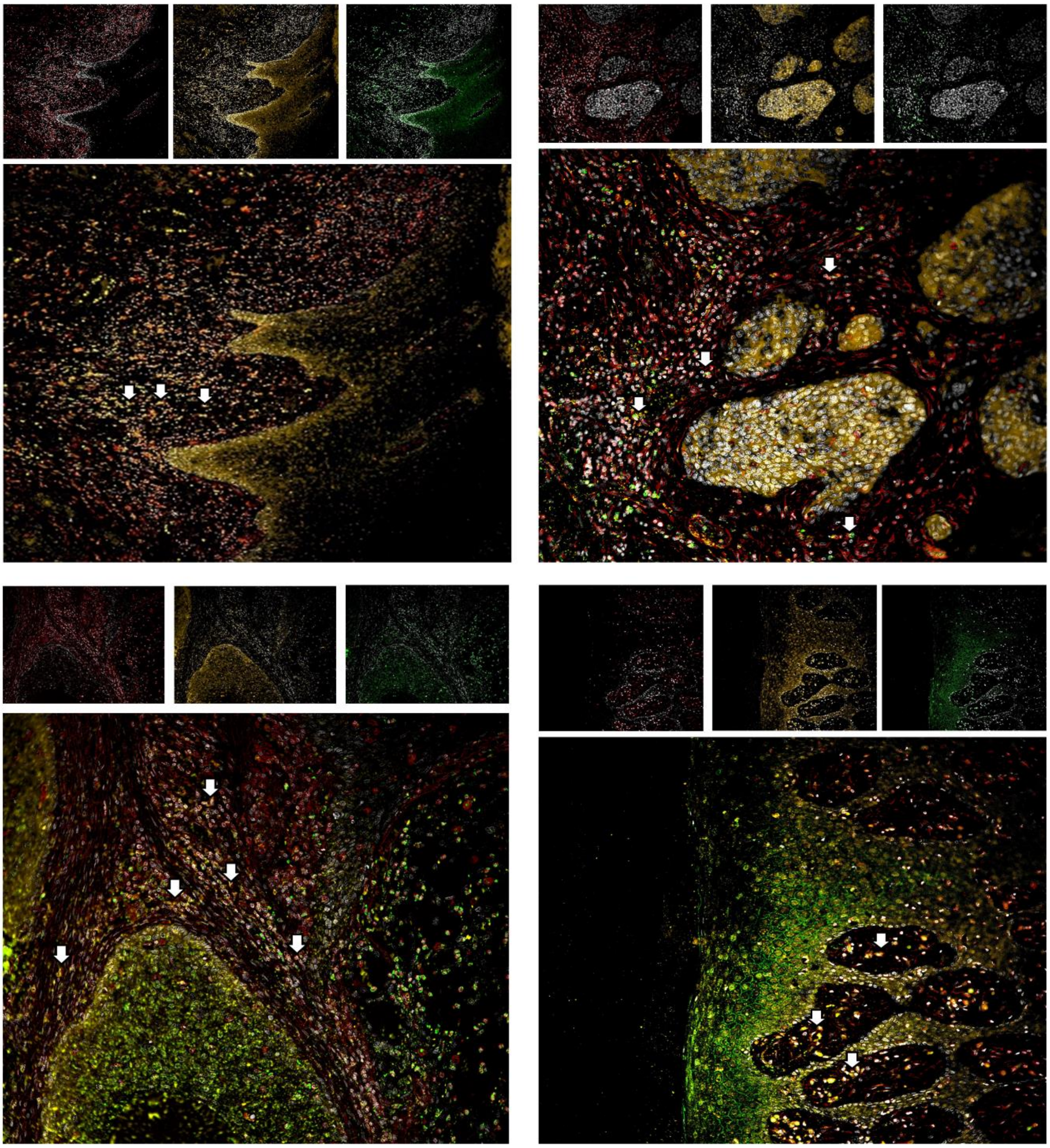

G Non-metastatic tumours

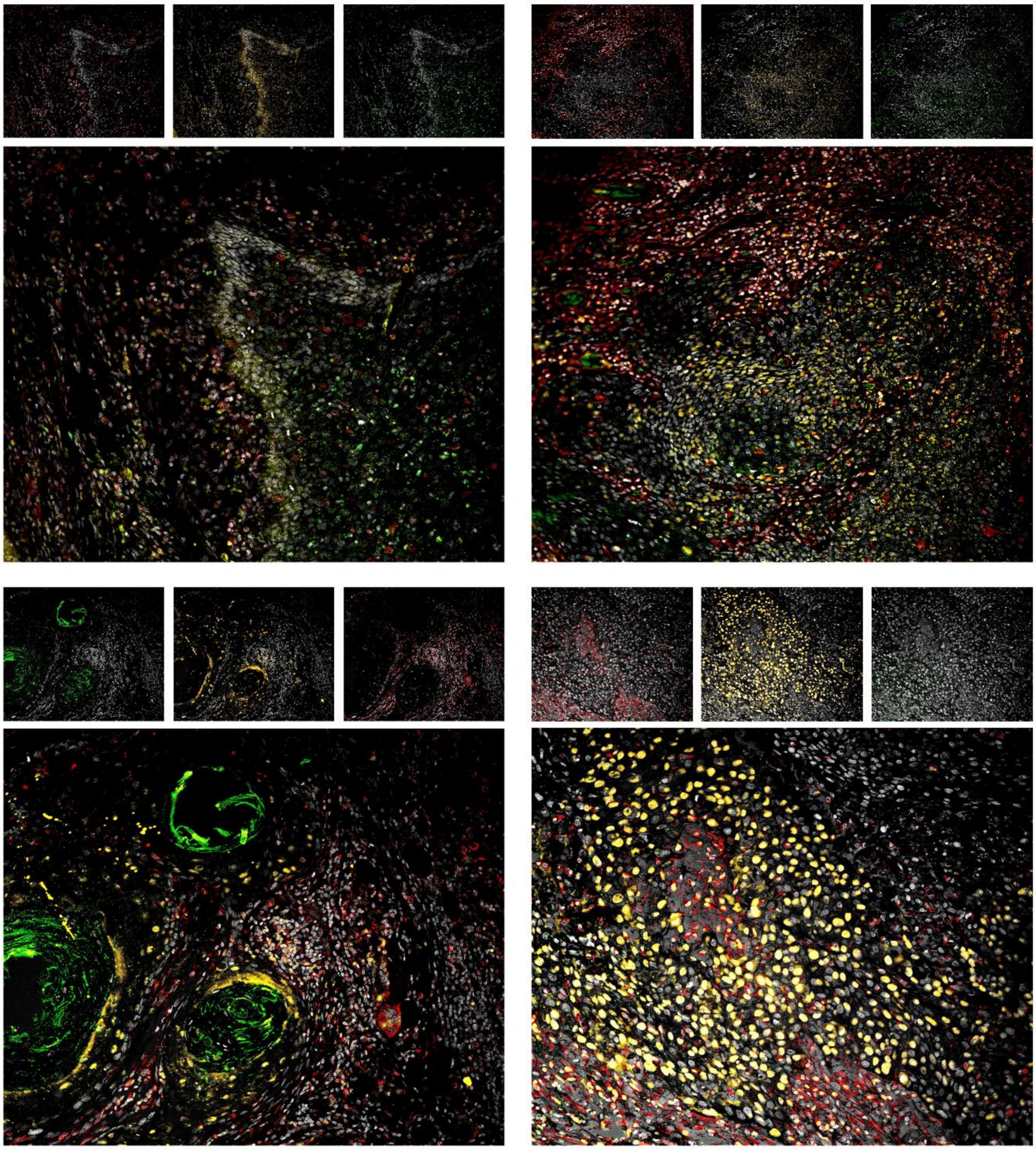


Figure S3 – Enrichment of EpCAM$^+$Vim$^+$CD24$^+$ cells in the stroma surrounding metastatic tumours in the second batch of specimens. **A,** Tiling of a stained and imaged slide into 20x fields of view and selection of a single field of view for segmentation. **B,** Segmentation into single nucleated cells using DAPI staining. **C,** Segmentation of cells in the stromal region using co-localisation of Vimentin and DAPI staining. **D,** This segmentation pipeline separates tumour body from stroma in image analysis. **E,** Cell identity measurements in the second batch of specimens. Quantification of the percentage of EpCAM$^+$Vim$^+$CD24$^+$ and EpCAM$^+$Vim$^+$CD24$^-$ cells in normal region (epithelium distant from the tumour), tumour body, and stromal region from metastatic and non-metastatic tumours. Recorded as the total percentage across the entire manually curated selection from the batch (tumour-stroma interface and normal region fields of view), rather than the average percentage per field of view (as shown in Fig. 2F for the first batch). **F-G,** The metastatic tumour fields (F) and non-metastatic tumour fields (G) from Fig. 2G, shown with separate channels at the top and the merge below. The separate channels are Vimentin (left, red), EpCAM (centre, yellow) and CD24 (right, green). All include DAPI nuclear stain. In the merge, white arrows highlight individual EpCAM$^{+}$Vim$^{+}$CD24$^{+}$ cells.

**Supplementary Figure S4**

A

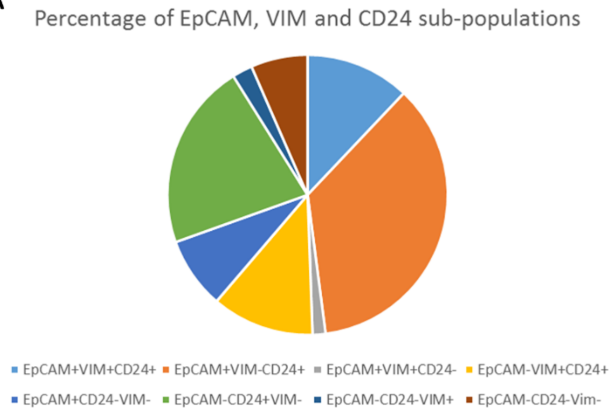


| Cancer Sub-populations in the Puram et al study | N | % |
|---|---|---|
| EpCAM+VIM+CD24+ | 267 | 12 |
| EpCAM+VIM-CD24+ | 795 | 36 |
| EpCAM+VIM+CD24- | 33 | 1 |
| EpCAM-VIM+CD24+ | 262 | 12 |
| EpCAM+CD24-VIM- | 183 | 8 |
| EpCAM-CD24+VIM- | 478 | 22 |
| EpCAM-CD24-VIM+ | 52 | 2 |
| EpCAM-CD24-Vim- | 145 | 7 |

B

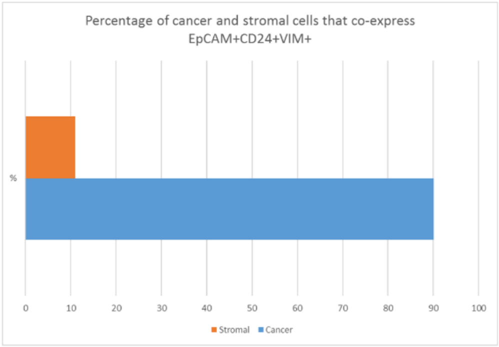


| Puram et al study | Cancer | Stromal |
|---|---|---|
| EpCAM+CD24+Vim | 267 | 29 |
| % | 90 | 11 |

Figure S4 – Analysis of EpCAM, CD24 and Vimentin expression in a published head and neck cancer scRNAseq dataset (Puram et al., 2017). **A,** Percentage of the cancer cells (excluding non-cancer cells) expressing each possible combination of EpCAM, CD24 and Vimentin, shown as a pie chart (left) and table (right). **B,** The percentage of EpCAM$^{+}$Vim$^{+}$CD24$^{+}$ cells within the whole dataset that are annotated as cancer (blue bar on bar chart) and non-cancer/stromal (orange bar on bar chart), shown as a bar chart (left) and table (right).

Puram, S. V., Tirosh, I., Parikh, A. S., Patel, A. P., Yizhak, K., Gillespie, S., Rodman, C., Luo, C. L., Mroz, E. A., Emerick, K. S., *et al.* (2017). Single-Cell Transcriptomic Analysis of Primary and Metastatic Tumor Ecosystems in Head and Neck Cancer. Cell *171*, 1611-1624 e1624.
